# FasL blockade suppresses liver metastasis and prolongs survival in small-cell lung cancer

**DOI:** 10.64898/2026.11.09.765391

**Authors:** Julia Saggau, Debora Bonasera, Anne Berghaus, Britta Jenke, Venelin Nikolov, Umut Akgül, Alexandra Florin, Reinhard Büttner, Filippo Beleggia, Maria Cartolano, H. Christian Reinhardt, Antonella Montinaro, Eva Rieser, Gianmaria Liccardi, Henning Walczak

## Abstract

Small-cell lung cancer (SCLC) is an aggressive malignancy characterised by early metastatic dissemination, frequent relapse and poor long-term survival. Despite advances in immunotherapy, durable disease control remains challenging, highlighting the need to identify therapeutic vulnerabilities beyond conventional tumour-cell killing and immune checkpoint inhibition. Here, we uncover an unexpected contribution of Fas-Fas ligand (FasL) signalling to SCLC metastatic progression and demonstrate the therapeutic potential of targeting this pathway. Using an autochthonous *Rb1/Trp53*-deficient mouse model, we show that genetic Fas deletion markedly suppresses liver metastatic disease, identifying a previously unrecognised contribution of Fas signalling to metastatic progression. Notably, despite this antimetastatic effect, Fas deletion during tumour initiation did not improve survival and was accompanied by pronounced alterations in pulmonary tissue architecture. As these alterations may reflect adverse consequences of Fas deletion in non-malignant pulmonary populations, we reasoned that pharmacological FasL blockade in established disease could provide therapeutic benefit while circumventing the detrimental consequences for pulmonary tissue homeostasis associated with genetic Fas deletion. Remarkably, only six administrations of a soluble Fas-Fc fusion protein over three weeks significantly prolonged survival and substantially reduced the incidence and burden of liver metastases. FasL blockade was also associated with enhanced inflammatory and immune-response transcriptional signatures in the liver, while its combination with anti-PD-1 increased pulmonary immune-cell accumulation without conferring additional survival benefit.

These data suggest that the biological consequences of targeting this pathway may depend on the timing and cellular context of intervention. Together, our findings identify Fas-FasL signalling as an unexpected regulator of SCLC metastatic progression and establish FasL blockade as a promising therapeutic strategy capable of prolonging survival in established disease beyond conventional immune checkpoint inhibition.

## Introduction

Small-cell lung cancer (SCLC) is a highly aggressive neuroendocrine malignancy characterised by rapid tumour growth, early metastatic dissemination and frequent relapse following treatment. Although SCLC initially responds to platinum-based chemotherapy, often combined with radiotherapy or immune checkpoint inhibition, the development of therapeutic resistance remains a major obstacle to durable disease control^1–3^. Genomic profiling has established near-universal inactivation of the tumour suppressors TP53 and RB1 as a defining molecular feature of SCLC^4^, providing the basis for genetically engineered mouse models in which conditional deletion of RB1 and TP53 recapitulates key features of human disease^5^. More recently, longitudinal genomic analyses have revealed extensive therapy-associated clonal diversification and tumour evolution, providing important insights into the mechanisms underlying disease recurrence^4^. Together, these findings emphasise the need to identify additional vulnerabilities that can be therapeutically exploited to improve disease control.

Beyond its characteristic genetic alterations, SCLC exhibits considerable transcriptional, metabolic and immunological heterogeneity^6,7^. Molecular classification has identified distinct cellular states associated with neuroendocrine differentiation, inflammatory signalling and differential therapeutic susceptibility^8^. Importantly, these states also exhibit divergent sensitivities to regulated cell death. Neuroendocrine and non-neuroendocrine SCLC populations display distinct dependencies on antioxidant defence mechanisms and susceptibility to ferroptosis^7^, while more recent evidence has demonstrated that metabolic programmes determine whether SCLC cells undergo apoptosis or ferroptosis in response to cellular stress^9^. More broadly, metabolic regulation has been shown to influence death-receptor-mediated apoptosis, with glucose availability and metabolic stress affecting signalling induced via Fas (CD95) and by TRAIL^10,11^. Moreover, the machinery controlling cell death can influence tumour development independently of its conventional role in eliminating malignant cells. Genetic ablation of Caspase-8 enhances tumour progression through MLKL-dependent necroptosis, inflammatory remodelling and acquisition of neuronal progenitor-like features^12^. These observations highlight a complex relationship between cell-death regulation, cellular plasticity and tumour progression in SCLC.

Death receptor-ligand systems of the tumour necrosis factor (TNF) and TNF receptor (TNFR) superfamilies (SFs) are important regulators of cellular survival, inflammation and tissue homeostasis^13–15^. Among these, the Fas-Fas ligand (FasL/CD95L) system is best characterised for its ability to induce extrinsic apoptosis^16,17^ by promoting recruitment of the adaptor FADD and procaspase-8, leading to formation of the death-inducing signalling complex (DISC) and initiation of the apoptotic cascade^13^. However, Fas signalling is not restricted to apoptosis. Depending on cellular context and signalling competence, Fas activation can also engage non-apoptotic pathways associated with inflammatory gene expression, cell proliferation, migration and invasion^13,17–19^. Indeed, Fas activation has been shown to induce the production of cytokines and chemokines that promote phagocyte recruitment, demonstrating that death-receptor engagement can actively shape inflammatory responses beyond the induction of apoptosis^20,21^. These non-apoptotic activities are particularly relevant in cancer, where resistance to Fas-mediated apoptosis does not necessarily abolish the capacity of tumour cells to respond to FasL. In apoptosis-resistant tumour cells, FasL stimulation was shown to enhance the migration and invasiveness of apoptosis-resistant tumour cells through activation of NF-κB and MAPK signalling^22^. Furthermore, FAS signalling promotes glioblastoma invasion through activation of the Yes-PI3K pathway^23^, while genetic deletion of Fas suppresses tumour formation and growth in ovarian and liver cancer models through mechanisms involving JNK-c-Jun signalling^24^.

The broader capacity of death receptors to promote inflammatory and tumour-supportive signalling is further supported by studies of the related TRAIL-TRAIL receptor (TRAIL-R) system. These have identified a non-enzymatic function of Caspase-8 as a scaffold for the assembly of FADD- and RIPK1-containing signalling complexes that promote NF-κB-dependent inflammatory gene expression^25^ and demonstrated that TRAIL-R signalling can promote tumour progression, metastatic dissemination and the establishment of an immunosuppressive microenvironment^26–28^. Thus, Fas and related death receptors can promote malignant progression under conditions in which their apoptotic activity is ineffective.

Importantly, the Fas-FasL system also regulates interactions between malignant cells and the immune microenvironment^14,17^. Although FasL expressed by cytotoxic lymphocytes can contribute to the elimination of susceptible target cells, FasL expression within tumours can conversely restrict antitumour immunity by promoting the apoptosis of activated effector lymphocytes. In particular, FasL expression by tumour-associated endothelial cells has been shown to establish a selective barrier against CD8⁺ T-cell infiltration^29^. Similarly, cancer-associated fibroblasts can suppress antitumour immunity through antigen-dependent elimination of CD8⁺ T cells involving FasL and PD-L2 signalling^30^. Furthermore, FasL-expressing myeloid-derived suppressor cells have been implicated in the elimination of tumour-infiltrating lymphocytes and resistance to immunotherapy, with FasL blockade improving therapeutic responses in preclinical tumour models^31^. These findings suggest that the consequences of targeting FasL may depend on the cellular compartment in which Fas signalling operates.

Together, these observations raise the possibility that Fas-FasL signalling, traditionally associated with the elimination of malignant cells, may instead promote metastatic progression while simultaneously compromising antitumour immunity. This duality is particularly relevant to SCLC, a malignancy characterised by early metastatic dissemination and limited durable responses to immune checkpoint blockade, in which both tumour-cell-intrinsic resistance and impaired immune surveillance contribute to therapeutic failure. Importantly, the biological consequences of targeting FasL cannot be predicted solely from its canonical apoptotic function, as blockade of this pathway could interfere with both tumour-promoting signalling and the elimination of antitumour lymphocytes. Yet, investigations of Fas signalling in SCLC have largely focused on chemotherapy-induced apoptosis and drug resistance³², leaving its contribution to metastatic progression, immune checkpoint responsiveness and tumour-immune interactions insufficiently understood.

Here, we investigated the contribution of Fas-FasL signalling to SCLC progression and its potential as a therapeutic target. Using an autochthonous mouse model initiated by adenoviral Cre-mediated deletion of Rb1 and Trp53, we demonstrate that concomitant genetic Fas deletion markedly suppresses liver metastases without improving survival, whereas pharmacological FasL blockade in established disease substantially reduces metastatic burden and significantly prolongs survival following a short treatment course. Our findings uncover an unexpected role for Fas-FasL signalling in SCLC metastatic progression and identify FasL blockade as a therapeutic strategy capable of improving disease outcome, either alone or in combination with PD-1-targeting immune checkpoint blockade (ICB). By integrating analyses of tumour progression, metastatic burden, immune-cell composition and transcriptional responses, we establish the biological and therapeutic relevance of the Fas-FasL axis in SCLC and provide a rationale for targeting this pathway beyond conventional ICB.

## Results

### Genetic ablation of Fas suppresses liver metastatic disease without improving survival in SCLC-bearing mice

Fas-FasL signalling has been extensively implicated in the regulation of apoptosis and immune homeostasis^13,32,33^, yet whether it contributes to SCLC progression and/or therapeutic response remains unknown. Consistent with the relative resistance of SCLC to Fas-mediated apoptosis^7,34–36^, several murine SCLC cell lines maintained high viability following stimulation with increasing concentrations of recombinant isoleucine zipper-tagged FasL (iz-FasL) (Extended Data Fig. 1a). Similarly, human SCLC cell lines exhibited heterogeneous responses to FasL stimulation, with several displaying considerable resistance compared with the FasL-sensitive non-small-cell lung cancer (NSCLC) cell line H460 (Extended Data Fig. 1b).

To assess whether Fas contributed to SCLC disease onset and progression we used an autochthonous mouse model of SCLC based on conditional deletion of *Rb1* and *Trp53* (referred to as RP mice)^5,37^. In this model, intratracheal administration of an adenovirus expressing Cre recombinase (Ad-CMV-Cre) induces recombination of the floxed alleles in infected pulmonary cells, resulting in the development of multifocal SCLC lesions over approximately four to six months (Fig. 1b). We then generated RP mice in which Fas could be conditionally depleted in parallel to Rb1 and p53 (*Rb1^fl/fl^*;*Trp53^fl/fl^*;*Fas^fl/fl^*mice), hereafter referred to as RPF. Intratracheal administration of Ad-CMV-Cre to the lungs of these mice permitted concurrent targeting of the three loci in infected cells which includes, SCLC-initiating cells (Fig. 1a), importantly, without being limited to them^38^. To assess tumour development, mice were monitored by magnetic resonance imaging (MRI). Approximately five months after viral administration, established tumour-bearing animals were included in the comparative analysis when they reached 2.5 mm^3^ as cumulative tumour burden. Interestingly, examination of RP mice (n=31) and RPF mice (n=13) showed that Fas deletion provided no difference in survival when compared, to RP controls (Fig. 1b). RPF mice showed a median survival of 55 days after tumour detection whilst RP mice showed a median survival of 52 days. Relative lung and liver weights remained comparable between the two experimental groups, whereas RPF mice exhibited increased relative spleen weight (Fig. 1c). Macroscopic and histological examination confirmed the presence of pulmonary SCLC lesions in the cohorts presenting similar tumour burden (Fig. 1d). Notably, however, more detailed histopathological assessment revealed pronounced alterations in pulmonary tissue architecture in RPF mice. Compared with RP controls, RPF lungs exhibited extensive disruption of normal alveolar organisation, with dense cellular regions and reduced recognisable airspaces (Fig. 1d-e). A similar architectural phenotype, characterised by dense cellular regions associated with extensive immune-cell infiltration, was observed in lungs lacking Fas independently of Rb1 and Trp53 deletion^39^ (Fig. 1f). These observations raise the possibility that the pulmonary alterations in RPF mice are not exclusively attributable to tumour progression but may also reflect the consequences of Fas deficiency in non-malignant pulmonary populations. Indeed, intratracheal Ad-CMV-Cre delivery does not restrict recombination to tumour-initiating cells and may additionally target other pulmonary cell populations, including resident immune cells. Consequently, Fas deletion in these compartments could contribute to inflammatory tissue remodelling and increased pulmonary cellularity, potentially confounding the histological assessment of tumour burden and influencing disease outcome independently of tumour growth.

**Figure 1:**
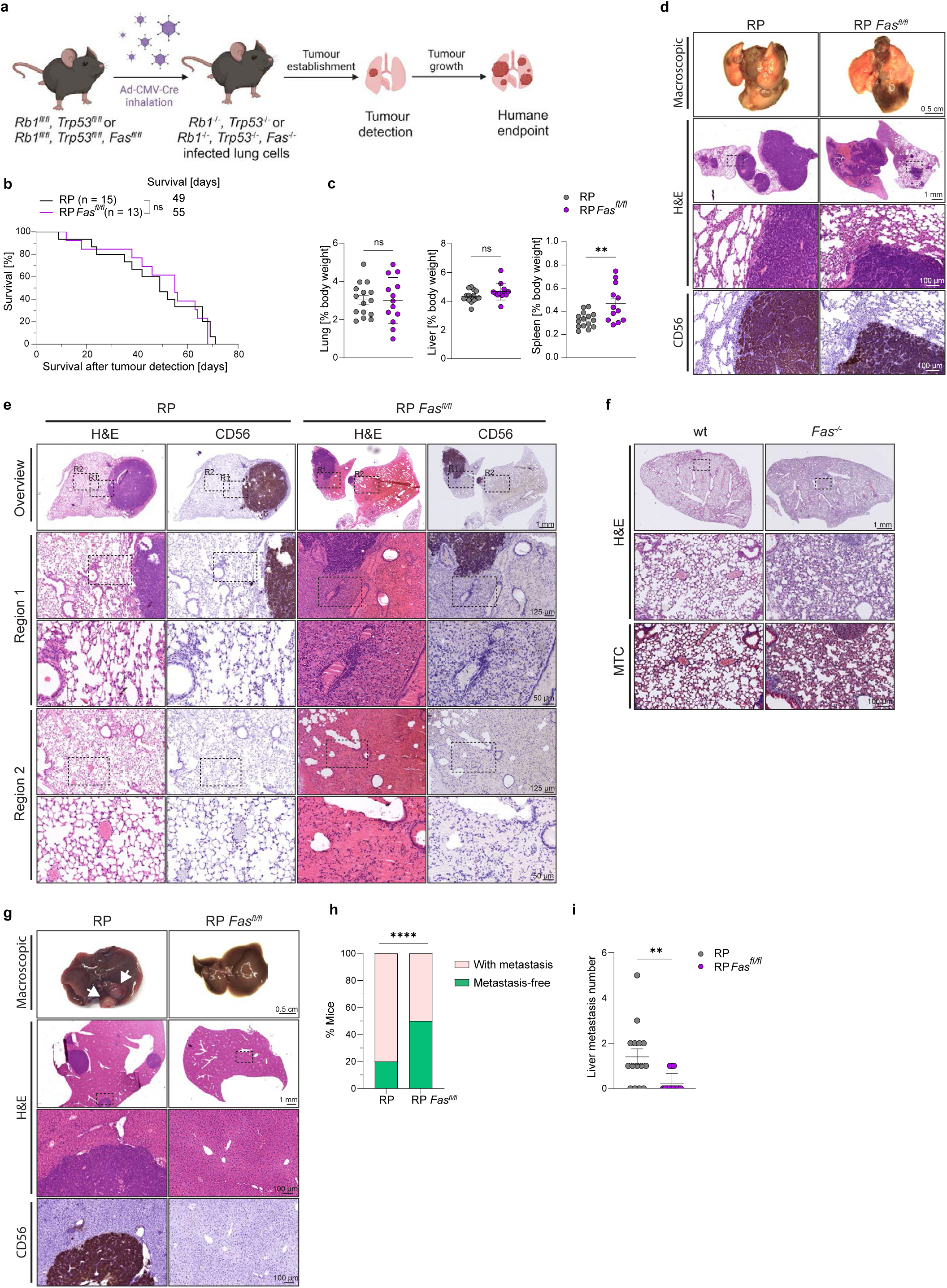
Tumour-specific deletion of Fas reduces metastatic burden of RP mice. **a**, Illustration of the experimental workflow for tumour induction and detection of lung tumours in RP and RP *Fas^fl/fl^* mice. Created with BioRender.com **b**, Kaplan-Meier survival analysis of RP and RP *Fas^fl/fl^* mice. Overall survival was assessed upon tumour detection, median post-treatment survival was determined via GraphPad Prism. Statistical analysis was performed via Log-rank (Mantel-Cox) test. **c**, Organ-to-body weight ratios for lung, liver and spleen measured at the humane endpoint. **d**, Representative images of lung tumour burden at the humane endpoint in RP and RP *Fas^fl/fl^* mice, shown as macroscopic images of the lungs, H&E staining and immunohistochemical staining for CD56. Boxed regions in the H&E overviews are shown at higher magnification in the row below. Scale bars: 0.5 cm (macroscopic), 1 mm (H&E overview), 100 µm (H&E and CD56 insets). **e**, Representative H&E and CD56 staining of lung tumour-adjacent tissue, including higher-magnification views of selected regions. Scale bars: 1 mm (overview), 125 µm (Region 1&2 overview) and 50 µm (higher magnification). **f**, Representative H&E and Masson’s trichrome (MTC) stainings of *Fas^-/-^*lungs at the humane endpoint compared to wt controls. Boxed regions in the H&E overviews are shown at higher magnification in the row below. Scale bars: 1 mm (H&E overview), 100 µm (H&E and CD56 insets). **g**, Representative images of liver metastatic burden at the humane endpoint, shown as macroscopic images of the livers, H&E staining and immunohistochemical staining for CD56. Arrows point at liver metastases. Boxed regions in the H&E overviews are shown at higher magnification in the row below. Scale bars: 0.5 cm (macroscopic), 1 mm (H&E overview), 100 µm (H&E and CD56 insets). **h**, Frequency of mice presenting with liver metastases or remaining metastasis-free at the humane endpoint as confirmed via MRI and immunohistochemical analysis. Statistical analysis was performed via two-tailed unpaired t-test with \*\*\*\**p* < 0.0001. **i**, Number of liver metastases quantified by MRI. All data present mean ± SEM. Statistical analysis was performed via two-tailed unpaired t-test with \**p* < 0.05.

Surprisingly, despite the absence of a survival advantage, genetic Fas deletion markedly reduced liver metastatic disease. Macroscopic examination revealed fewer liver lesions in RPF mice compared to RP (Fig. 1g). H&E staining and CD56 immunohistochemistry confirmed reduced liver tumour involvement. Consistently, RPF mice exhibited a significantly increased proportion of metastasis-free animals compared with RP controls (Fig. 1h), together with significant reductions in liver metastatic number (Fig. 1i).

Together, these findings identify an unexpected contribution of Fas signalling to liver metastatic progression in SCLC. Although genetic Fas deletion markedly reduced metastatic burden, this effect was not accompanied by improved survival. Importantly, the pronounced pulmonary architectural alterations observed in RPF mice, which resemble those associated with Fas deficiency independently of tumourigenesis, suggest that loss of Fas in non-malignant pulmonary populations may contribute to disease severity and potentially obscure any survival benefit associated with reduced metastatic burden. Thus, while our findings implicate Fas signalling in SCLC metastatic progression, the broader consequences of genetic Fas deletion highlight the difficulty of distinguishing its effects on tumour biology from those on pulmonary tissue homeostasis.

### Pharmacological inhibition of FasL prolongs survival in an autochthonous mouse model of small-cell lung cancer

Given the marked reduction in liver metastases and the pronounced pulmonary architectural alterations associated with genetic Fas deletion, we next investigated whether pharmacological FasL blockade could provide therapeutic benefit in established SCLC while avoiding the potential consequences of receptor deletion during tumour initiation on pulmonary tissue homeostasis. This approach also allowed us to assess the consequences of targeting Fas-FasL signalling without permanently deleting the receptor during tumour initiation, while providing a more translationally relevant strategy. To this end, we employed a murine Fas-Fc fusion protein (mFas-Fc), comprising the extracellular domain of murine Fas fused to the Fc region of mouse IgG2a, which functions as a soluble decoy receptor to neutralise FasL^40^ (Fig. 2a).

**Figure 2:**
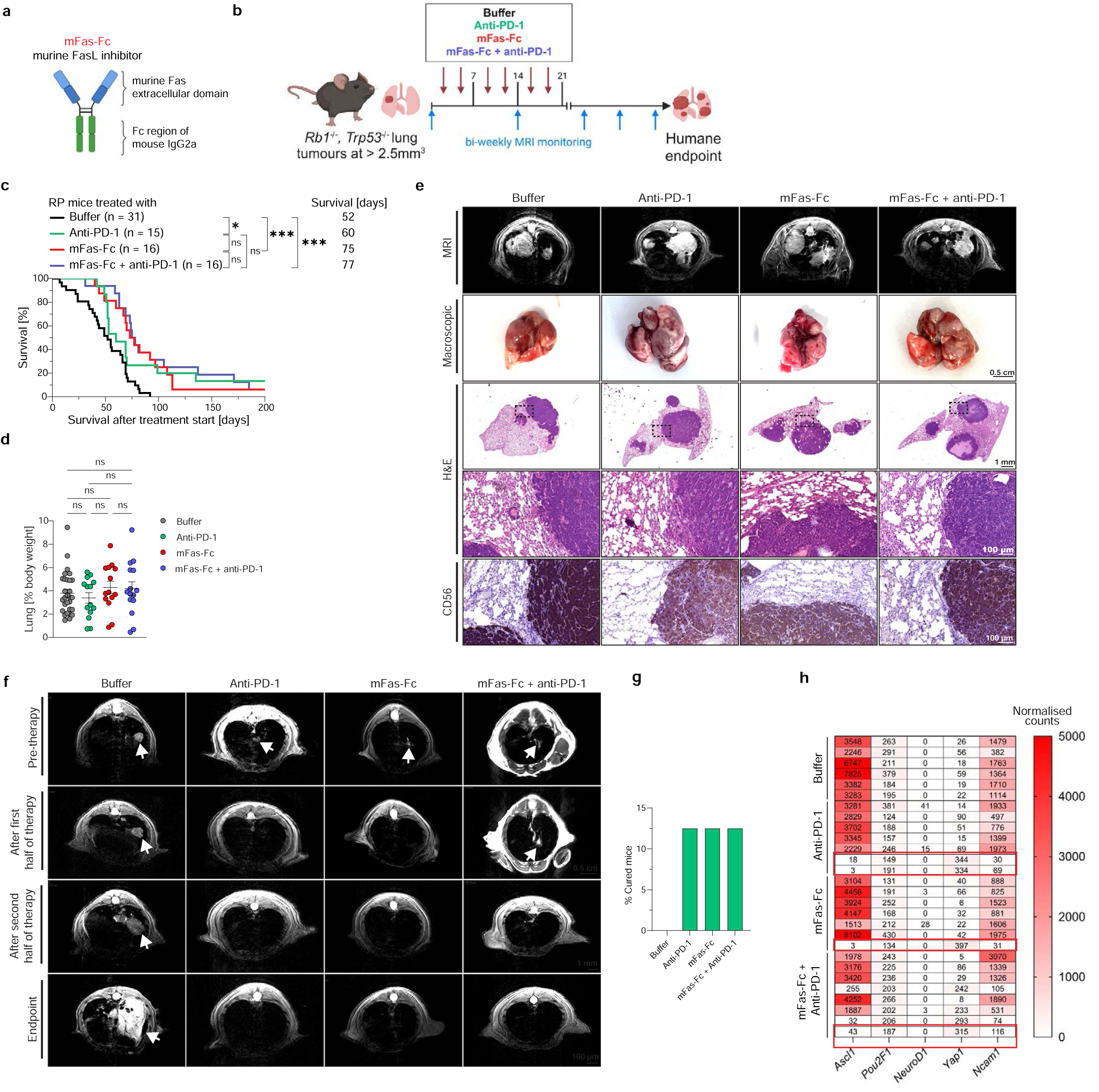
The biotherapeutic FasL inhibitor mFas-Fc significantly prolongs survival in RP mice. **a**, Schematic representation of the murine FasL inhibitor mFas-Fc. Created with BioRender.com **b**, Illustration of the experimental workflow including tumour induction, detection and treatment of lung tumours in RP mice. Created with BioRender.com **c**, Kaplan-Meier survival analysis of RP mice treated with Buffer, Anti-PD1, mFas-Fc and the combination. Overall survival was assessed upon treatment start, median post-treatment survival was determined via GraphPad Prism. Statistical analysis was performed via Log-rank (Mantel-Cox) test with \**p* < 0.05 and \*\*\**p* < 0.001. **d**, Percentage of lung size relative to total body weight measured at the humane endpoint. **e**, Representative images of lung tumour burden at the humane endpoint in mice treated with Buffer, anti-PD-1, mFas-Fc, or the combination, shown as MRI, macroscopic images of the lungs, H&E staining and immunohistochemical staining for CD56. Boxed regions in the H&E overviews are shown at higher magnification in the row below. Scale bars: 0.5 cm (macroscopic), 1 mm (H&E overview), 100 µm (H&E and CD56 insets). **f**, Representative MRI scans of cured mice at initial tumour detection before therapy start and in the treatment course. Arrows point at lung tumours. **g**, Frequency of mice with complete lung tumour regression. **h**, 3’ RNA-sequencing analysis for SCLC subtype markers in the lung tissue at the humane endpoint. Gene expression changes are shown as normalized counts and colour-coded in red. Mice with complete tumour regression are highlighted by red boxes.

To assess the therapeutic efficacy of FasL inhibition against established disease, tumour development was monitored by MRI as explained above. Following confirmation of pulmonary tumour formation, mice were randomised into four experimental groups receiving vehicle (n = 31), anti-PD-1 (n = 15), mFas-Fc (n = 16), or combined mFas-Fc and anti-PD-1 (n = 16). The experimental design was intended to determine whether blockade of FasL could improve the outcome of established SCLC, either as monotherapy or by enhancing the therapeutic activity of PD-1 blockade. Following treatment initiation, pulmonary tumour progression was assessed longitudinally by MRI at two-week intervals, and animals were monitored for overall survival until the experimental endpoint. Treatments were administered twice weekly for three consecutive weeks, corresponding to six administrations per animal (Fig. 2b). The treatment duration was selected to minimise potential immune responses against the rat-derived anti-PD-1 antibody, and an identical schedule was applied across all experimental groups.

Survival analysis revealed a significant therapeutic benefit following FasL inhibition (Fig. 2c). Vehicle-treated RP mice exhibited a reported survival of 52 days after treatment initiation, whereas anti-PD-1 monotherapy extended survival to 60 days. Notably, treatment with mFas-Fc increased survival to 75 days, while the combination of mFas-Fc and anti-PD-1 resulted in a survival of 77 days. Although both mFas-Fc-containing regimens significantly improved survival compared with vehicle, the combination did not confer a statistically significant additional advantage over mFas-Fc monotherapy. These findings demonstrate that pharmacological blockade of FasL can substantially prolong survival in mice with established SCLC.

Surprisingly, the observed survival benefit was not accompanied by a corresponding reduction in pulmonary tumour burden. Relative lung weights remained comparable across the four experimental groups (Fig. 2d), while representative MRI, macroscopic examination and histological analysis demonstrated the persistence of substantial pulmonary tumour lesions following treatment (Fig. 2e). Haematoxylin and eosin (H&E) staining and immunohistochemical detection of the neuroendocrine marker CD56 confirmed the presence of comparable SCLC lesions in the examined lungs. Consistent with these observations, longitudinal MRI-based assessment revealed continued growth of pulmonary tumours in most animals, without a consistent treatment-associated reduction in primary tumour volume (Extended Data Fig. 2a-c). Notably, although body weights remained comparable across treatment groups, mice receiving combined mFas-Fc and anti-PD-1 treatment exhibited significantly increased relative spleen weights (Extended Data Fig. 2d, e), indicative of an enhanced systemic immune activation.

Despite the limited overall effect on pulmonary tumour burden, individual treated mice exhibited striking therapeutic responses. Serial MRI analyses performed before treatment, during the treatment course, following treatment completion and at the experimental endpoint revealed progressive regression of initially detectable pulmonary lesions in a subset of animals (Fig. 2f). Such responses were observed in mice receiving anti-PD-1, mFas-Fc or the combination, with the analysis identifying a small proportion of animals in each treated cohort showing apparent complete tumour regression (Fig. 2g). Transcritpional assessment of SCLC-associated lineage markers^8^, including *Ascl1*, *Pou2f3*, *Neurod1*, *Yap1* and *Ncam1*, provided further evidence supporting the absence of detectable SCLC-associated molecular signatures in selected responding animals (Fig. 2h).

Collectively, these findings establish that FasL blockade substantially improves survival in autochthonous SCLC despite having limited effects on primary pulmonary tumour burden, suggesting that its therapeutic activity involves processes beyond direct suppression of primary tumour growth.

### FasL blockade in combination with anti-PD-1 promotes immune-cell accumulation in the SCLC-bearing lung

Because Fas-FasL signalling has a central role in regulating immune-cell survival, we next investigated whether the therapeutic effect of mFas-Fc was associated with changes in the immune compartment of SCLC-bearing mice. To this end, we established a separate experimental cohort of RP mice in which pulmonary tumours were induced by intratracheal administration of Ad-CMV-Cre, as described above. Five months after tumour induction, mice with established pulmonary tumours were randomised to receive vehicle, anti-PD-1, mFas-Fc or the combination. Treatments were administered twice weekly for three consecutive weeks. At the completion of this treatment period, mice were sacrificed, and lungs and spleens were collected for comprehensive flow cytometric characterisation of lymphoid and myeloid immune-cell populations (Fig. 3a). Three weeks after treatment initiation, neither body weight nor relative lung and spleen weights differed between treatment groups (Fig. 3b-d).

**Figure 3:**
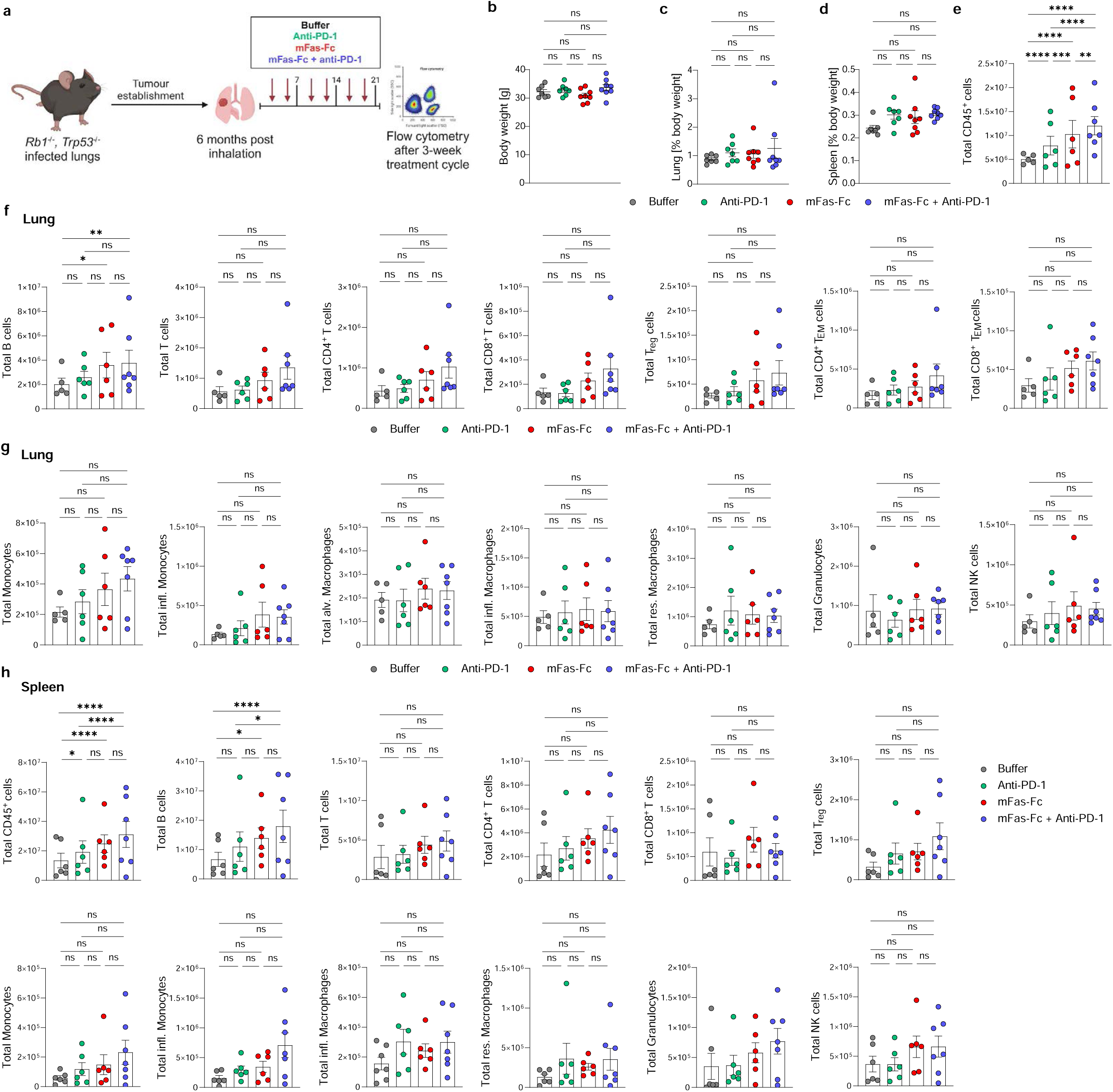
Combination of FasL blockade and anti-PD1 promotes immune cell infiltration in the lungs of RP mice. **a**, Schematic illustration of the experimental setup for flow cytometric analysis of immune cells in the lungs and spleens of treated RP mice. Created with BioRender.com **b**, Body weight after the three-week treatment cycle in RP mice treated with Buffer, anti-PD-1, mFas-Fc or the combination. **c,d**, Percentage of lung (**c**) and spleen (**d**) size relative to total body weight after three-week treatment cycle. **e**, Total number of CD45⁺ immune cells in the lung after the three-week treatment cycle in mice treated with Buffer, anti-PD-1, mFas-Fc or the combination. **f,g**, Total numbers of lymphocytes (**f**) and myeloid cells (**g**) in the lung after the three-week treatment cycle. **h**, Flow cytometric analysis of immune cell subsets in the spleens of RP mice following a three-week treatment cycle. All populations represent the total number of immune cells per organ. Statistical analysis was performed via two-way ANOVA using GraphPad Prism with \**p* < 0.05, \*\**p* < 0.01 and \*\*\*\**p* < 0.0001.

Quantification of pulmonary CD45-positive leukocytes revealed a pronounced increase in immune-cell accumulation in mice treated with mFas-Fc and anti-PD-1, while monotherapy with either agent was associated with more modest changes (Fig 3e). Importantly, the immune-cell increase following the combination treatment occurred at a defined treatment endpoint and was not accompanied by a corresponding reduction in overall pulmonary tumour burden, suggesting that treatment-associated immune-cell accumulation can occur independently of substantial primary tumour regression.

To determine whether this effect reflected alterations in the immune-cell make-up, we next quantified the abundance of specific lymphoid and myeloid populations (Fig. 3f-g). Among the populations examined, B cells exhibited a significant increase following combination therapy compared with the relevant control groups (Fig. 3f). Total T cells, including CD4- and CD8-positive subsets, also tended to increase following combined treatment, although these differences did not reach statistical significance (Fig. 3f). Similarly, regulatory T cells (Tregs) and monocytes did not exhibit consistent statistically significant differences across the experimental groups (Fig. 3g).

Additional characterisation of pulmonary immune populations, including effector-memory CD4-positive and CD8-positive T cells, alveolar and interstitial macrophages, inflammatory monocytes, granulocytes and natural killer (NK) cells, did not reveal substantial treatment-associated differences (Fig. 3g). Thus, the increase in total pulmonary leukocyte abundance was not accompanied by a broad, statistically significant expansion of individual lymphoid or myeloid populations.

We next determined the effect of FasL inhibition on systemic immune-cell abundance. To do so, we examined splenic populations collected at the same three-week experimental endpoint (Fig. 3h) Combined treatment resulted in increased numbers of total CD45-positive leukocytes and B cells in the spleen, whereas most other immune-cell populations remained largely unchanged (Fig. 3h). The gating strategies used to identify the respective lymphoid and myeloid populations are provided in Extended Data Fig. 3.

Together, these findings demonstrate that combined FasL/PD-1 inhibition promotes immune-cell accumulation within tumour-bearing lungs and is associated with alterations in systemic immune-cell abundance. However, the absence of consistent changes in individual effector-cell populations suggests that additional mechanisms may contribute to the survival benefit observed following FasL blockade.

### FasL inhibition suppresses liver metastatic disease and is associated with enhanced inflammatory and immune-response signatures

The observation that FasL inhibition prolonged survival without substantially affecting primary pulmonary tumour burden prompted us to investigate whether the treatment influenced disease progression at secondary anatomical sites. Since the RP model develops liver metastases, mirroring one of the most common sites of metastatic dissemination in human SCLC patients, we examined liver involvement across the four treatment groups.

Although relative liver weights were comparable across experimental groups (Fig. 4a), treatment significantly altered the incidence of liver metastases. Compared with vehicle-treated controls, a substantially greater proportion of animals receiving anti-PD-1, mFas-Fc or the combination remained free of detectable liver metastases (Fig. 4b). Furthermore, quantitative MRI analysis demonstrated significant reductions in the number and total volume of liver metastatic lesions following treatment (Fig. 4c).

**Figure 4:**
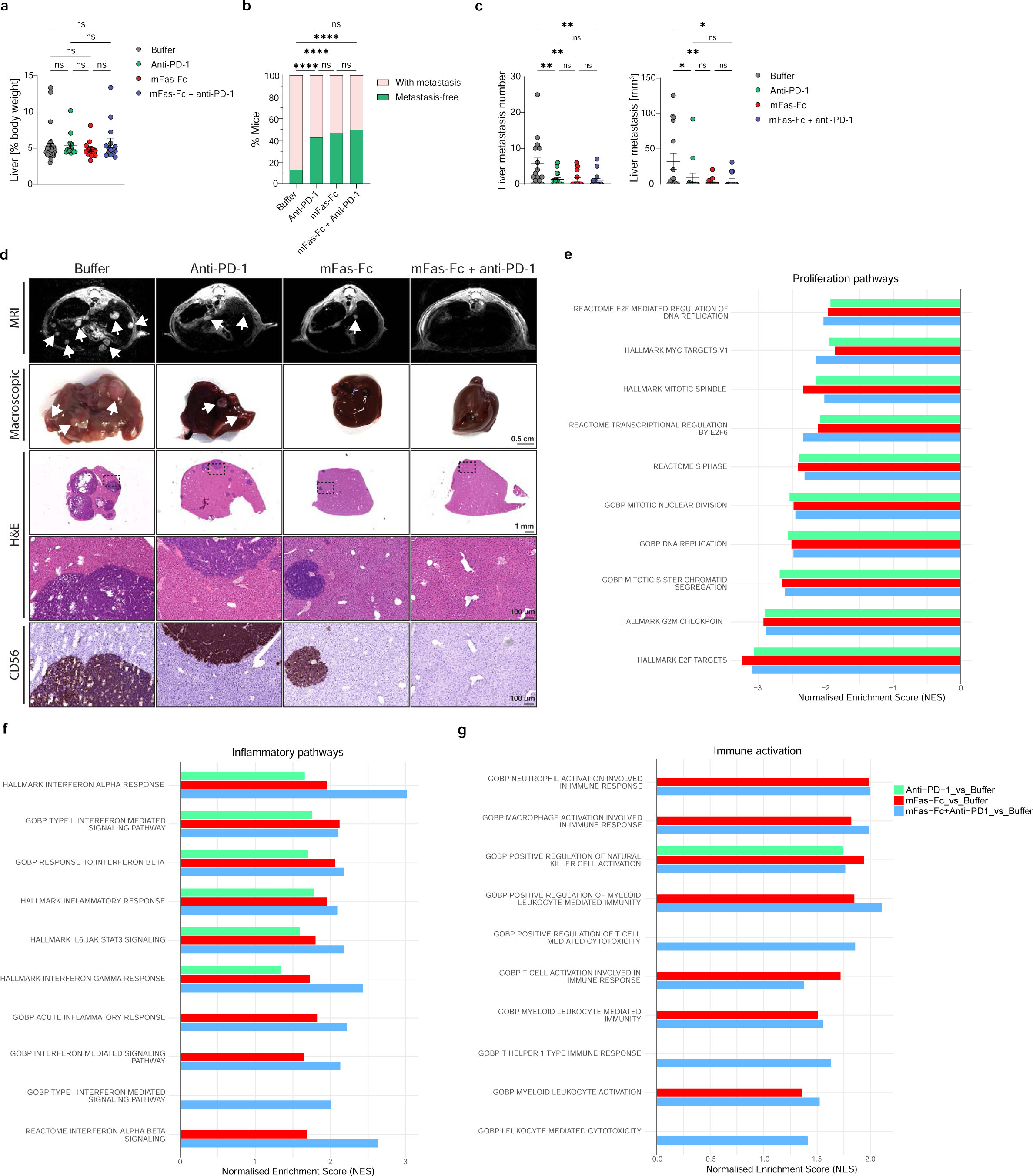
FasL blockade reduces incidence and size of liver metastases in RP mice. **a**, Percentage of liver size relative to total body weight measured at the humane endpoint. **b**, Frequency of mice presenting with liver metastases or remaining metastasis-free at the humane endpoint as confirmed via MRI and immunohistochemical analysis. **c**, Number and cumulative volume of liver metastases quantified by MRI. For each mouse, the total metastatic volume represents the sum of all liver metastatic nodules. All data present mean ± SEM. Statistical analysis was performed via two-tailed unpaired t-test with \**p* < 0.05, \*\**p* < 0.01, \*\*\**p* < 0.001, \*\*\*\**p* < 0.0001. **d**, Representative images of liver metastatic burden at the humane endpoint in mice treated with Buffer, anti-PD-1, mFas-Fc, or the combination, shown as MRI, macroscopic images of the livers, H&E staining and immunohistochemical staining for CD56. Arrows point at liver metastases. Boxed regions in the H&E overviews are shown at higher magnification in the row below. Scale bars: 0.5 cm (macroscopic), 1 mm (H&E overview), 100 µm (H&E and CD56 insets). **e-g**, Gene set enrichment analysis of 3’ RNA-sequencing data from liver tissue at the humane endpoint, showing normalised enrichment scores (NES) for gene sets related to proliferation (e), inflammatory signalling (f) and immune activation (g). Scores are shown for each treatment relative to Buffer as indicated. Only gene sets with an adjusted *p* value < 0.05 are plotted.

These observations were corroborated by macroscopic and histopathological examination. Vehicle-treated animals frequently exhibited multiple liver lesions, whereas treated mice displayed considerably less extensive metastatic involvement. H&E staining and immunohistochemical detection of CD56 confirmed the marked reduction in liver SCLC lesions in the cohorts that received mFas-Fc and/or anti-PD1 treatment (Fig. 4d).

To investigate molecular changes associated with the reduction in metastatic disease, we analysed transcriptional programmes in liver samples from the different treatment groups. Gene set enrichment analysis (GSEA) revealed negative enrichment of proliferation-associated programmes, including E2F and MYC targets, DNA replication, mitotic spindle organisation and G2/M checkpoint signatures (Fig. 4e). These findings were accompanied by pronounced enrichment of inflammatory-response pathways, particularly following mFas-Fc treatment and its combination with anti-PD-1 (Fig. 4f). Enriched signatures included type I and type II interferon (IFN-I/II) responses, interleukin 6-JAK-STAT3 signalling and broader inflammatory programmes.

Consistent with these findings, analysis of immune-response-associated pathways revealed enrichment of transcriptional signatures linked to macrophage, neutrophil and NK cell activation, as well as T-cell activation, T-helper 1 responses and leukocyte-mediated cytotoxicity (Fig. 4g). These effects were particularly prominent in the mFas-Fc-receiving treatment groups.

Collectively, these findings identify suppression of liver metastatic disease as a prominent consequence of FasL inhibition in autochthonous SCLC. Importantly, this effect occurs despite the persistence of substantial primary pulmonary tumour burden and is accompanied by enhanced inflammatory and immune-response transcriptional activity in the liver.

## Discussion

The limited success of current treatments for SCLC underscores the need to identify therapeutic vulnerabilities beyond conventional tumour-cell killing and immune checkpoint inhibition. Here, we identify FasL-induced signalling as a therapeutically actionable regulator of SCLC progression. Using an autochthonous mouse model of established disease^5^, we demonstrate that pharmacological FasL blockade significantly prolongs survival and markedly reduces liver metastatic burden, despite having limited effects on primary pulmonary tumour growth. Importantly, genetic depletion of Fas during tumour initiation similarly suppressed liver metastases but failed to reproduce the survival benefit observed following systemic FasL blockade. These findings reveal a previously unknown contribution of Fas-FasL signalling to SCLC progression and suggest that metastatic dissemination and overall disease outcome are influenced by distinct, context-dependent functions of this pathway.

A particularly striking observation is the effect of FasL blockade on liver metastases compared with primary pulmonary tumour burden. Although mFas-Fc treatment prolonged survival from 52 to 75 days after treatment initiation, pulmonary tumours generally continued to progress. Instead, FasL blockade significantly reduced the incidence, number and size of liver metastatic lesions. This observation is particularly relevant in light of the immunological properties of the liver, where metastatic tumours can exploit mechanisms of peripheral immune tolerance^41,42^. It has been demonstrated that liver metastases recruit FasL-expressing monocyte-derived macrophages capable of inducing apoptosis of activated Fas-positive, tumour-specific CD8⁺ T cells, thereby compromising systemic antitumour immunity and limiting responsiveness to immune checkpoint inhibition^42^. Our findings raise the possibility that FasL blockade interferes with similar immunosuppressive interactions in the liver metastatic microenvironment of SCLC. Consistent with this hypothesis, transcriptional profiling revealed enrichment of IFN-I- and IFN-II-associated, inflammatory and immune-response programmes in the livers of mFas-Fc-treated mice. Whether these changes reflect preservation of antitumour lymphocytes, altered myeloid-cell function or differences in metastatic tumour content remains to be established. We have previously shown that upregulation of these transcriptional programmes reflects inflammatory cell death priming^43^, raising the possibility that such mechanisms contribute to the observed antimetastatic effects. However, we did not detect corresponding differences in cell-death-associated transcriptional signatures or immunohistochemical markers. This may reflect the transient and spatially heterogeneous nature of cell-death events, which could occur at earlier stages of metastatic dissemination, including during circulation or initial liver colonisation, rather than within established metastatic lesions analysed at the experimental endpoint.

The potential involvement of tissue-specific Fas-FasL signalling is especially intriguing because this pathway can exert opposing effects on metastatic progression depending on the cellular and anatomical context. Whereas FasL expression by tumour-associated endothelial and myeloid populations can promote immune evasion through elimination of cytotoxic lymphocytes^29,31^, Fas-dependent apoptosis can also restrict the survival of disseminated malignant cells. In osteosarcoma, FasL expression within the pulmonary microenvironment contributes to the elimination of Fas-expressing tumour cells and limits metastatic colonisation^44^. More recently, it was demonstrated that glucocorticoid receptor activation promotes metastatic seeding by enabling disseminated tumour cells to evade Fas-dependent immune cytotoxicity^45^. These findings illustrate the opposing biological consequences of Fas-FasL signalling depending on whether the principal responding population comprises malignant cells or components of the antitumour immune response. In SCLC, where several cell lines exhibit limited sensitivity to FasL-induced apoptosis, inhibition of FasL may therefore preferentially interfere with non-apoptotic, tumour-supportive or immunosuppressive functions of the pathway rather than protect malignant cells from apoptotic elimination.

Our genetic experiments further reveal the complexity of Fas signalling during SCLC development. Although Fas deletion markedly reduced liver metastatic burden, it did not improve survival and was accompanied by pronounced alterations in pulmonary tissue architecture. These observations suggest that the consequences of genetic *Fas* deficiency cannot be exclusively attributed to tumour-cell-intrinsic mechanisms. Importantly, the intratracheal Ad-CMV-Cre system used to initiate SCLC induces recombination of the floxed *Rb1*, *Trp53* and *Fas* alleles in susceptible infected cells, without restricting recombination exclusively to the malignant epithelial compartment. Indeed, it has previously been demonstrated that intratracheal Ad-CMV-Cre can transduce alveolar macrophages in genetically engineered lung cancer models, resulting in recombination and altered macrophage behaviour^38^. Consequently, although the same viral intervention targets all three conditional alleles, the cellular populations undergoing recombination and the efficiency of deletion at each locus remain important considerations. In the absence of cell-type-resolved recombination analyses, the altered pulmonary architecture observed in RPF mice cannot be unequivocally attributed to changes in tumour growth or tumour-cell differentiation.

This distinction is particularly important given the established role of Fas in maintaining immune-cell homeostasis and pulmonary tissue integrity. Germline *Fas* deficiency causes aberrant lymphocyte accumulation, lymphadenopathy and splenomegaly, accompanied by lymphocytic infiltration of peripheral tissues, including the lungs and liver^46,47^. Furthermore, selective deletion of Fas in T cells can induce profound disturbances in lymphocyte homeostasis, inflammatory pulmonary infiltration and progressive fibrosis, demonstrating that disruption of Fas signalling in immune cells alone can profoundly alter lung pathology^48^. Fas signalling also contributes to the resolution of pulmonary inflammation by promoting the elimination of recruited inflammatory macrophages, providing an additional mechanism through which impaired Fas activity may influence lung architecture independently of tumourigenesis^49^. In agreement with the importance of death-receptor signalling in pulmonary pathology, we recently demonstrated that Fas contributes to disease progression in a genetically defined model of spontaneous lung fibrosis driven by enhanced LUBAC signalling^39^. Although these disease models differ substantially, they emphasise that Fas signalling influences pulmonary tissue homeostasis through mechanisms extending beyond malignant cells. It is therefore conceivable that the altered lung morphology and survival phenotype observed in RPF mice arise, at least in part, from changes in immune-cell homeostasis or tissue organisation associated with Fas deficiency, rather than exclusively from altered tumour-cell-intrinsic mechanisms. Whether the RPF phenotype involves local immune-cell expansion, inflammatory remodelling or other consequences of Fas loss remains unresolved.

The distinction between genetic *Fas* deletion and pharmacological FasL blockade is further reinforced by the different experimental timing and anatomical scope of these interventions. Whereas genetic recombination occurs during tumour initiation and may affect several pulmonary cell populations, systemic mFas-Fc treatment is initiated after the establishment of detectable disease and can interfere with FasL-dependent interactions across multiple tissues. The latter may be particularly relevant in liver metastases, where FasL-expressing immune populations can contribute to local and systemic immune suppression^42^. Consistent with an immunomodulatory effect, combined FasL and PD-1 blockade increased pulmonary CD45⁺ leukocyte accumulation and B-cell abundance, while liver transcriptional analyses revealed enrichment of inflammatory and immune-response pathways. However, these changes were not accompanied by consistent increases in individual effector T-cell populations, and the enhanced inflammatory signatures cannot be interpreted as direct evidence of restored antitumour immunity. Nevertheless, the convergence of reduced liver metastases, increased immune-associated transcriptional activity and improved survival supports further investigation of FasL-dependent regulation of the metastatic microenvironment.

Importantly, six administrations of mFas-Fc over three weeks were sufficient to significantly prolong survival and substantially reduce liver metastatic burden in mice with established SCLC. The treatment schedule was deliberately restricted to accommodate the concomitant administration of a rat-derived anti-PD-1 antibody and minimise the potential development of host immune responses against the heterologous antibody following repeated administration. To ensure comparability across experimental groups, the same treatment schedule was applied to mFas-Fc monotherapy, despite the use of a fully murine fusion protein. Importantly, the efficacy of sustained FasL blockade remains unexplored, and prolonged treatment with mFas-Fc could potentially provide additional therapeutic benefit. Determining whether sustained FasL blockade can further improve disease control will therefore be important, although the kinetics of tumour development and the logistical demands of this autochthonous model complicate the evaluation of extended treatment schedules.

The therapeutic implications of our findings are particularly relevant given the limited efficacy of immune checkpoint inhibition in SCLC. Although the addition of PD-L1 inhibitors to first-line platinum-etoposide chemotherapy has improved survival in extensive-stage SCLC, the magnitude and durability of these benefits remain limited^50,51^. Moreover, several PD-1-directed strategies have failed to meet their primary overall-survival objectives in phase III clinical trials, including pembrolizumab combined with first-line platinum-etoposide chemotherapy and nivolumab-based maintenance treatment following chemotherapy^52,53^. Although more recent therapeutic advances, including lurbinectedin combined with atezolizumab as maintenance treatment and DLL3-directed T-cell engagement with tarlatamab, have demonstrated improvements in clinical outcomes, durable disease control remains a substantial challenge^51,54^. Against this background, the ability of FasL blockade to prolong survival despite persistent primary pulmonary disease identifies a potentially complementary therapeutic strategy targeting aspects of SCLC progression that may be insufficiently controlled by conventional immune checkpoint inhibition. Importantly, although combined mFas-Fc and anti-PD-1 treatment enhanced pulmonary immune-cell accumulation, it did not confer a significant additional survival advantage over mFas-Fc monotherapy. These findings therefore do not establish synergy between FasL and PD-1 blockade, but instead indicate that FasL inhibition can independently provide therapeutic benefit in a setting where PD-1 blockade alone has limited efficacy.

It remains to be established which cellular compartments and molecular mechanisms are responsible for the therapeutic activity of FasL blockade in SCLC. In particular, the cellular sources of FasL within primary and metastatic lesions, the contribution of Fas-dependent immune-cell apoptosis and the consequences of genetic Fas deletion across different pulmonary cell populations require further investigation. Moreover, whether the reduction in liver metastatic burden reflects impaired dissemination, reduced metastatic colonisation or restricted growth of established metastases cannot be resolved from the current experiments. Defining these mechanisms will be essential to establish the contribution of tumour-intrinsic and microenvironmental Fas signalling to SCLC progression and to determine how FasL blockade might be incorporated into future therapeutic strategies.

Our findings establish Fas-FasL signalling as an unexpected regulator of metastatic progression in SCLC, extending its biological significance beyond the canonical induction of tumour-cell apoptosis. Importantly, the ability of a short course of FasL blockade to markedly suppress liver metastases and prolong survival, despite continued primary tumour growth, demonstrates that therapeutically meaningful disease control can be achieved without substantial regression of the primary malignancy. These findings identify FasL blockade as a promising strategy for targeting metastatic disease in SCLC and provide a foundation for developing therapeutic approaches directed at FasL-dependent interactions between malignant cells and their microenvironment.

## Author contributions

Conceptualisation: H.W., G.L. Supervision: H.W., G.L.

Animal licenses and documentation: J.S., D.B., E.R.

Experimental design: J.S., D.B., A.B., H.W., G.L.

Preliminary experimental work: J.S., D.B., A.M., F.B.

In-vivo experiments: J.S., D.B., B.J., V.N., A.B.

In-vitro experiments: A.B., A.M.

IHC stainings: A.F., R.B. J.S., D.B.

Design and analysis of flow cytometry data: J.S., D.B., A.M.

RNA-seq analysis: U.A., M.C., D.B.

Data discussion: H.C.R.

Visualisation: D.B., J.S., A.B., G.L.

Writing: J.S., D.B., H.W., G.L.

## Acknowledgements

We thank all members of the Liccardi and Walczak laboratories as well as the members of the CRC1399 for their helpful discussions. We are grateful to the Histology Core Facility of SFB1399, the CECAD in vivo Research Facility at the University of Cologne, the animal facilities of the Medical Faculty at the University of Cologne, and the Cologne Center for Genomics for their technical support and assistance. We further thank Louisa Grauvogel (University of Cologne) for her support with initial RNA-seq data analyses. Finally, we thank Dirk Wiedermann (Max Planck Institute for Metabolism Research, Cologne), Holger Grüll and Thorsten Persigehl (University Hospital Cologne, University of Cologne) for their support with their respective MRI facilities.

This study was funded by the Deutsche Forschungsgemeinschaft (DFG) through SFB1399 (413326622) (H.C.R, M.C., F.B., G.L., H.W.), SFB1530 (455784452) (R.B., M.C., H.C.R., G.L., H.W.), SFB1403 (414786233) (H.W.), the Alexander von Humboldt Foundation (H.W.), a Wellcome Trust Investigator Award (214342/Z/18/Z to H.W.), a Medical Research Council Grant (MR/S00811X/1 to H.W.), a Cancer Research UK Programme Grant (A27323 to H.W.), the CANcer TARgeting (CANTAR) Network grant by the Ministry of Art and Science of the State of NRW, Germany (H.W., G.L.), the E.I. Stiftung Kölner Krebsforschung Grant (G.L.), the Center for Molecular Medicine Cologne (CMMC) Individual & Tandem Project Program through Project A07 (H.W.) and B04 (G.L.), the Institute of Biochemistry I, Medical Faculty, University of Cologne (G.L., H.W.).

## Availability of Data and Materials

All unique reagents generated in this study are available upon reasonable request from the lead contact.

## Materials and Methods

### Mice

The *Rb1^fl/fl^*, *Trp53^fl/fl^* (RP) mouse model was kindly provided by Anton Berns (NKI). *Fas^fl/fl^* mice were derived from an in-house breeding colony originally established with animals obtained from The Jackson Laboratory (Bar Harbor, ME, USA), as previously described^55^. All mice were kept on a C57BL/6 background. Experiments included both male and female mice, comprising a total of 102 RP and 13 RPF mice.

SCLC tumour induction was performed as previously described^5,37^. In brief, RP or RP *Fas^fl/fl^* mice received a single intratracheal administration of adenovirus encoding Cre recombinase under the control of the CMV promoter (Ad5CMVCre, WC-U of Iowa-5, University of Iowa) at 8-12 weeks of age (2.5 × 10⁷ PFU per mouse in PBS, 75 µl per instillation). Tumour establishment and progression was assessed by biweekly MRI starting at 5 months of age. Once tumours reached a volume of 2.5 mm³, mice were randomised to receive intraperitoneal (i.p.) injections of 200 µL vehicle (DPBS, 14190144, Thermo Fisher Scientific; n = 31), 250 µg anti-PD-1 (RMP1-14, BXC-BE0146-50MG, Biozol; n = 15), 250 µg mFas-Fc (WuXi Biologics; n = 16)^40^, or the combination of anti-PD-1 and mFas-Fc (n = 16). Treatments were administered twice weekly for three weeks, after which mice were aged until they reached humane endpoint criteria.

For the time-point analysis, mice were randomised into treatment groups at 6 months after tumour induction and treated for three weeks with vehicle (n = 5), anti-PD1 (n = 6), mFas-Fc (n = 6) or the combination thereof (n = 7). Mice were culled on completion of the treatment cycle and organs were harvested for further analyses, to assess early disease development.

The mice were housed in individually ventilated cages under controlled conditions, including a consistent 12-hour light/dark cycle, at the animal facilities of the CECAD Research Centre as well as the animal facilities of the Medical Faculty at the University of Cologne. They had unrestricted access to food and water and were maintained at stable room temperature and humidity levels. All mice were kept under specific pathogen-free (SPF) conditions, with pathogen screenings conducted quarterly. Animals were monitored daily and euthanized via cervical dislocation once they met pre-defined endpoint criteria. All experiments were conducted in accordance with German animal protection laws and were approved by the relevant institutional ethics committees and local government authorities (*Landesamt für Natur, Umwelt und Verbraucherschutz Nordrhein Westfalen*, approval number Az 81-02.04.2020.A101).

### MR Imaging

Magnet resonance images were performed to determine the lung tumour volume. For this, mice were anesthetised with 2,5% isoflurane and then scanned using the 9.4 T preclinical Bruker MR scanner (Bruker Corporation, Ettlingen, Germany) equipped with a small rodent coil. Overview scans were acquired by performing 5 dummy scans (TE: 2,3 ms; TR: 107,82 ms; Flip angle: 30°). Images of the abdomen were acquired with axial slice orientation (TE: 29 ms; TR: 2464.959 ms; slice thickness: 0.7 mm). The images were exported in DICOM format and spleen volumes were analysed using the HOROS analysis software. The results were graphed using GraphPad Prism 11.0.2.

### Tissue processing and Immunohistochemistry

For immunohistochemical analysis, organs were harvested and fixed in 4% paraformaldehyde for 24 h and subsequently embedded in paraffin. Lungs and livers were cut at 3 µM and subjected to H&E, CD56 (ZYT-RBK050, Biozol) or MTC (Sigma-Aldrich, HT15) staining. Image analysis was performed using QuPath.

### Flow Cytometry

For the flow cytometric analysis, mice were randomised into treatment groups at 6 months post-instillation and treated for three weeks as described in the in-vivo paragraph before. Mice were culled on completion of the treatment cycle, and lungs as well as spleens were harvested. Single-cell suspension preparation and the flow cytometry workflow were performed as previously described^39^. Samples were measured on a BD FACSymphony™ A3 Cell Analyser. Data were analysed using FlowJo™ v10.10 and GraphPad Prism 11.0.2. The gating strategy is illustrated in Extended Data Figure 4.

### RNA isolation and 3’ mRNA sequencing

Snap-frozen lung and liver tissue was homogenised in lysis buffer (Buffer RLT Plus supplemented with 40 µl of 1 M DTT per ml and 0.5% Reagent DX) with 5 mm stainless steel beads (69989, Qiagen, Hilden, Germany) in a TissueLyser II (85300, Qiagen). Total RNA was subsequently extracted with the RNeasy Plus Mini Kit (74136, Qiagen) following the manufacturer’s instructions. For 3’ mRNA sequencing, 2 µg of RNA was submitted to the Cologne Center for Genomics (CCG), where library preparation was carried out using the Lexogen QuantSeq 3’ mRNA sequencing protocol. Sequencing was performed on a NovaSeq 6000 platform using a single-read (SR) 1×100 bp sequencing protocol.

Downstream analyses were performed in R (v4.6.1). Raw gene-level counts were analysed with DESeq2^56^ (v1.52.0). Genes with fewer than 10 total counts were excluded, and counts were normalised using the DESeq2 median-of-ratios method. Each treatment group was compared with Buffer controls using the DESeq2 Wald test, and the resulting Wald statistic was used to rank genes for gene set enrichment analysis (GSEA). Ranked gene lists were analysed with fgsea^57^ (v1.38.0) using the Hallmark, GO Biological Process, Reactome and BioCarta collections from MSigDB^58^,retrieved via msigdbr (v26.1.1) with gene set size 10–500. Pathways with a Benjamini–Hochberg-adjusted *p*-value < 0.05 were considered significant. Normalised enrichment scores were plotted with ggplot2^59^ (v4.0.3).

### Cell culture

Murine SCLC cell lines were cultured in DMEM (PAN-Biotech, P04-03550) and human SCLC cell lines were cultured in in RPMI (PAN-Biotech, P04-16500), both supplemented with 10% FBS (Sigma-Aldrich, F7524) and 1% Penicillin/Streptomycin (PAN-Biotech, P06-07100). Cells were seeded in 96-well plates and treated with increasing concentrations of recombinant isoleucine zipper-tagged FasL (iz-FasL) as indicated in the respective figure. 48 h after treatment, a CellTiter-Glo® Luminescent Cell Viability Assay (Promega, G7573) was performed. Luminescence was measured in a Spark® Multimode Microplate Reader (TECAN).

### Statistical analysis

Data shown represent the mean ± SEM, as indicated in the figure legends. Statistical significance was determined using unpaired, two-tailed parametric Student’s t-test, Log-rank (Mantel-Cox) test, one- or two-way ANOVA with Tukey’s multiple comparisons test with \**p* < 0.05, \*\**p* < 0.01, \*\*\**p* < 0.001, \*\*\*\**p* < 0.0001. All statistical analyses were performed using GraphPad Prism 11.0.2.

## Supplementary Materials

**Table S1.** Antibodies for flow cytometric analysis of lungs and spleens.

| **Antibody** | **Source** | **Identifier** | **Dilution** |
|---|---|---|---|
| NK1.1 | BD Biosciences | Cat# 741715 | 1:200 |
| CD4 | BD Biosciences | Cat# 564667 | 1:200 |
| CD45 | BD Biosciences | Cat# 612924 | 1:250 |
| CD11b | BD Biosciences | Cat# 565080 | 1:250 |
| CD3 | BioLegend | Cat# 563565 | 1:200 |
| Ly6G | BD Biosciences | Cat# 741994 | 1:250 |
| CD62L | BioLegend | Cat# 104441 | 1:100 |
| CD68 | BioLegend | Cat# 137021 | 1:100 |
| CD279 | BD Biosciences | Cat# 744546 | 1:100 |
| Cd11c | BioLegend | Cat# 117334 | 1:100 |
| Ly6C | BioLegend | Cat# 128041 | 1:150 |
| CD16/32 | BD Biosciences | Cat# 553144 | 1:100 |
| CD44 | BD Biosciences | Cat# 745922 | 1:200 |
| CD206 | BioLegend | Cat# 141706 | 1:100 |
| CD152 | BD Biosciences | Cat# 564332 | 1:100 |
| CD95 | BD Biosciences | Cat# 557653 | 1:100 |
| Foxp3 | Thermo Fisher Scientific | Cat# 7-5773-82 | 1:100 |
| B220 | BioLegend | Cat# 103232 | 1:200 |
| CD8 | BD Biosciences | Cat# 612898 | 1:250 |
| CD95L | BioLegend | Cat# 106609 | 1:75 |

---

**Extended Data Figure 1:**
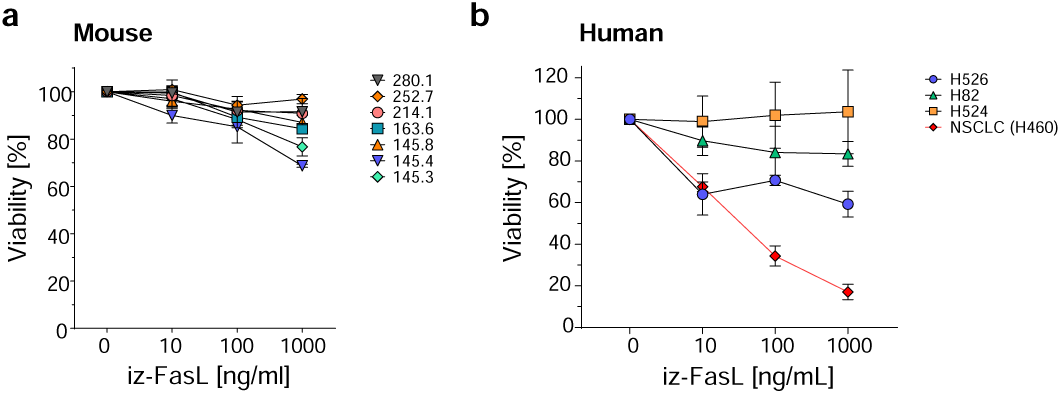
Human and murine SCLC cells are resistant to FasL-induced cell death. a, b, Murine and human SCLC cell lines were treated for 48 h with indicated concentrations of iz-FasL. Cell viability was measured via CellTiter-Glo assay. Data present mean ± SEM (n = 3 independent experiments per cell line).

**Extended Data Figure 2:**
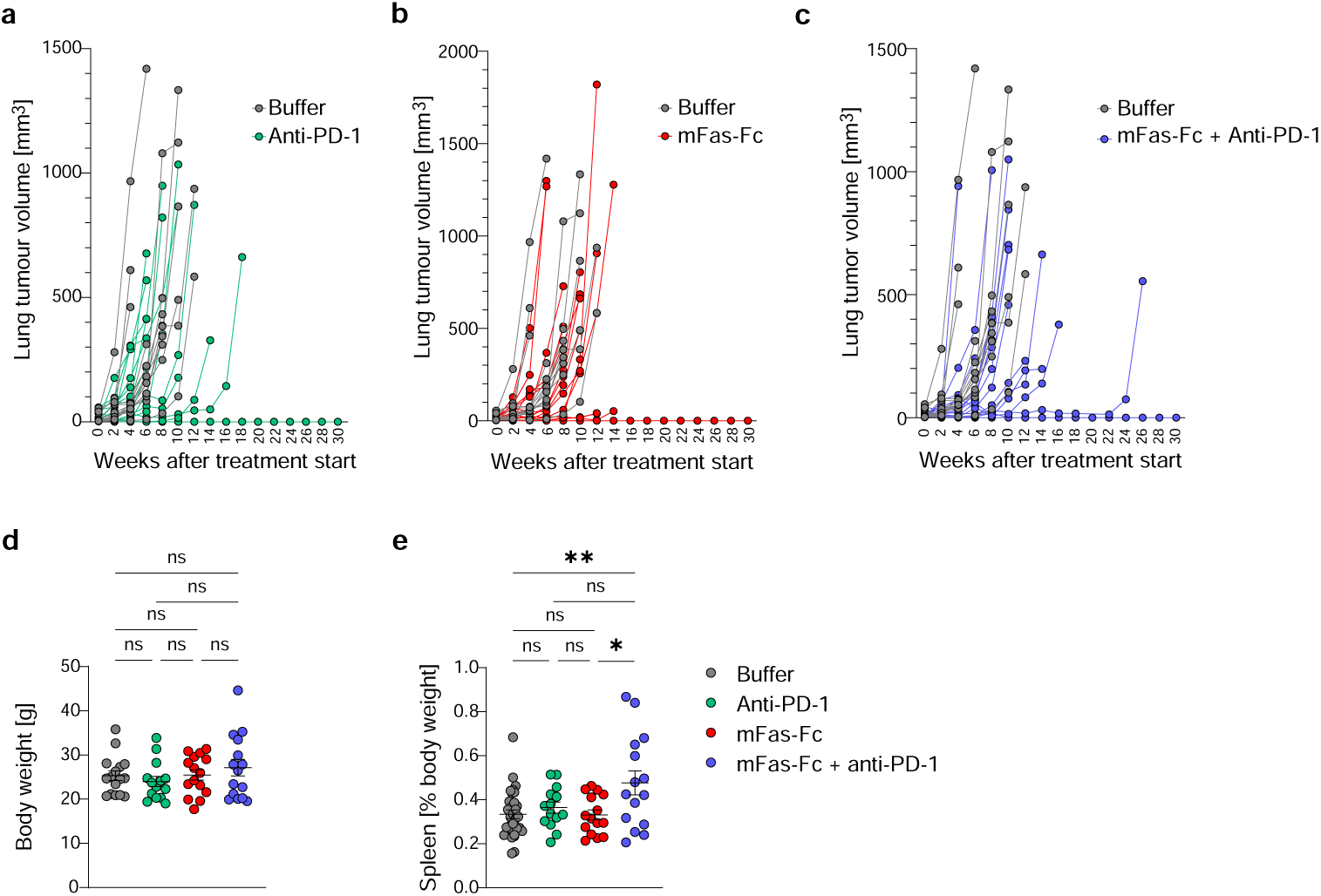
Treatment of RP mice with FasL blockade or anti-PD1 does not affect lung tumour progression. **a**, Body weight at the humane endpoint in RP mice treated with Buffer, anti-PD-1, mFas-Fc or the combination. **b**, Percentage of spleen size relative to total body weight measured at the humane endpoint. **c-e**, Total lung tumour volume of individual mice over the course of treatment, as determined by MRI, comparing Buffer-treated controls with anti-PD-1 (**c**), mFas-Fc (**d**) or the combination of mFas-Fc and anti-PD-1 (**e**). Each line represents one animal. Statistical significance was assessed via One-way ANOVA using GraphPad Prism with \**p* < 0.05 and \*\**p* < 0.01. Related to Figure 2.

**Extended Data Figure 3:**
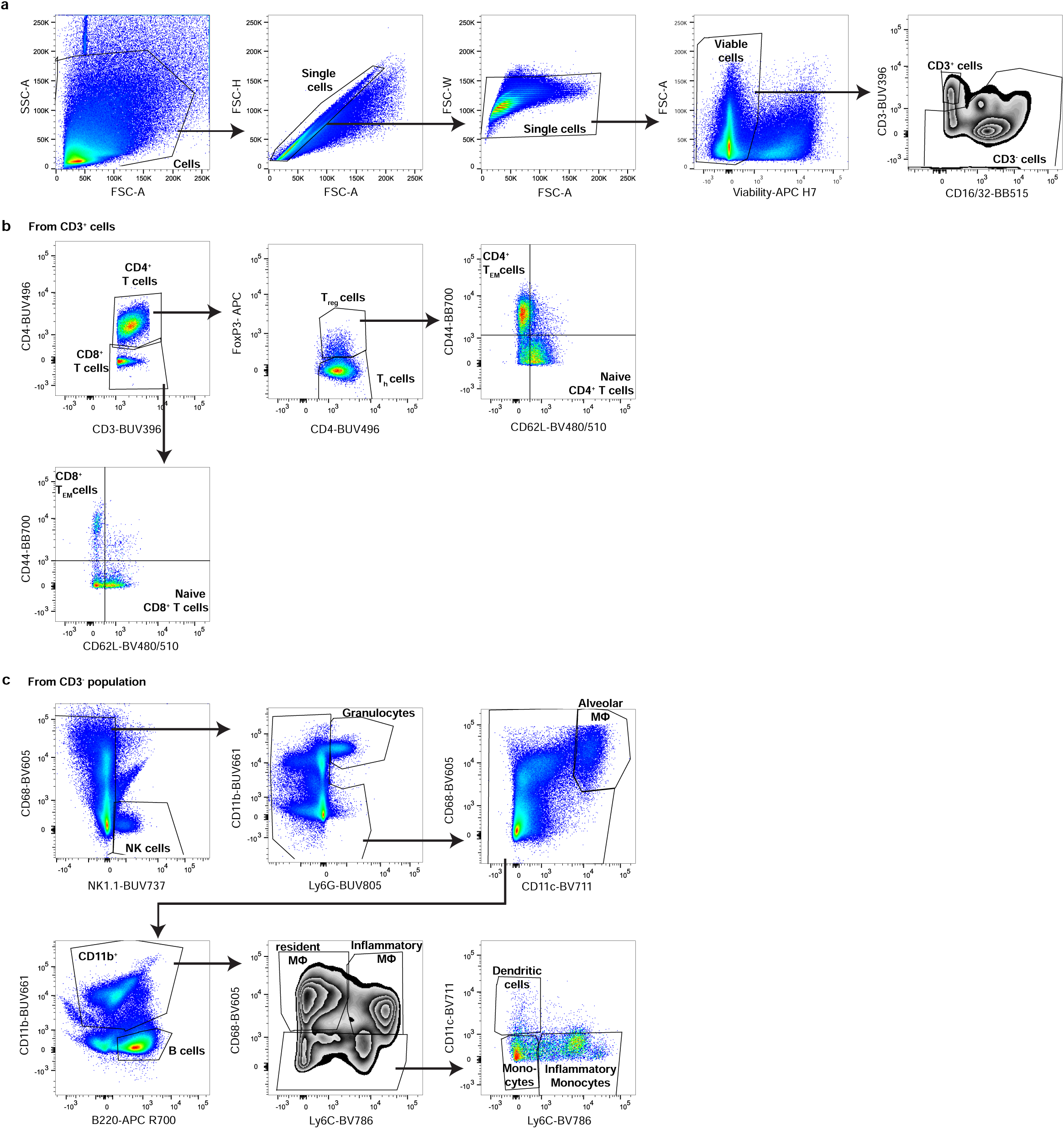
Gating strategy for identification of immune cell subsets in the lungs and spleens of RP mice. a, Flow cytometry gating strategies for identification of immune cells from mouse lungs. **b**, Identification of T-cell subsets from CD3^+^ immune cells. **c**, Gating of myeloid and B-cell populations from CD3^-^ immune cells. Related to Figure 3.

